# Measuring the Potential for Community Rescue Under Environmental Stress

**DOI:** 10.64898/2026.01.29.702686

**Authors:** Dong-Hao Zhou, Shuai Li, Liang Tang, Quan-Guo Zhang

## Abstract

The widely used half maximal inhibitory concentration and related indices, e.g., minimum inhibitory concentration and mutant prevention concentration, are *in vitro* measures of response of biological or biochemical processes to inhibitor substances. We propose to adapt the above concepts for developing standard measures of functional resistance and rescue of ecological communities under environmental pollution. First, a “half inhibitory concentration” index (IC_50_) is defined as the concentration at which a pollutant reduces a community-level function to 50% of the baseline state immediately after exposure; note that this is different from the “half maximal effective concentration” (EC_50_) index commonly used for estimating drug safety. Second, a “half community rescue prevention concentration” index (CRPC_50_) is the concentration at which a pollutant can prevent community-level ecological function from recovering to 50% of the baseline state after a prolonged period of recovery. CRPC_50_ is supposedly higher than IC_50_. We performed two experiments with natural aquatic microbial communities and considered respiration rate as a surrogate of whole-community function. The IC_50_ and CRPC_50_ of the herbicide glyphosate and the heavy metallic salt copper sulfate were estimated by dose-response curves. One specific finding is that CRPC_50_ values of glyphosate were typically higher than IC_50_ by > 2.4 folds, while for copper sulfate the CRPC_50_ values did not significantly differ from IC_50_. This suggested that our studied microbial communities showed little adaptive response to the latter pollutant agent. Standardized measures of functional resistance and community rescue potential such as those proposed in the present study would be comparable across environmental stress agents and study systems, and improve our study of short- and long-term consequences of environmental pollutants for microbial community functioning.

## INTRODUCTION

Dose-response relationships are prevalent in chemical and biological systems. While such response curves are often non-linear and need to be described by models with more than two parameters such as Michaelis-Menten equation, extracting a single parameter that is comparable among relationships is often desirable (Ankomah & Levin, 2012; Brain & Cousens, 1989; Cedergreen et al., 2005; Finney, 1979; Michaelis & Menten, 1913; van Ewijk & Hoekstra, 1993). For example, the half maximal inhibitory concentration (IC_50_) index is widely used to quantify the efficiency of a certain substance to inhibit a given chemical or biological process.

Antibacterial susceptibility assays typically involve comparing the efficacies of different drugs, or the susceptibility of different tested microbial strains. In such studies, the IC_50_ index is specifically defined as the concentration at which a drug inhibits the growth of a specific strain by 50% (Fig. 1a) (Mitosch & Bollenbach, 2014; Soothill et al., 1992). Commonly used was also the minimum inhibitory concentration (MIC) index, defined as the lowest concentration at which a drug blocks the growth of a given strain (Andrews, 2008; Hogg, 2013). Note that, in practice, the MIC measure may not be estimated from a dose-response curve; instead, an investigator may simply assign the lowest drug concentration used in a specific assay that blocks visible bacterial growth as MIC (Fig. 1a) (Barnes V et al., 2023; Blondeau & Fitch, 2019; Kadeřábková et al., 2024). Not surprisingly, antibacterial testing work is often concerned with MIC rather than IC_50_, as complete blocking of bacterial growth is pursued in antibacterial development.

**Figure 1.**
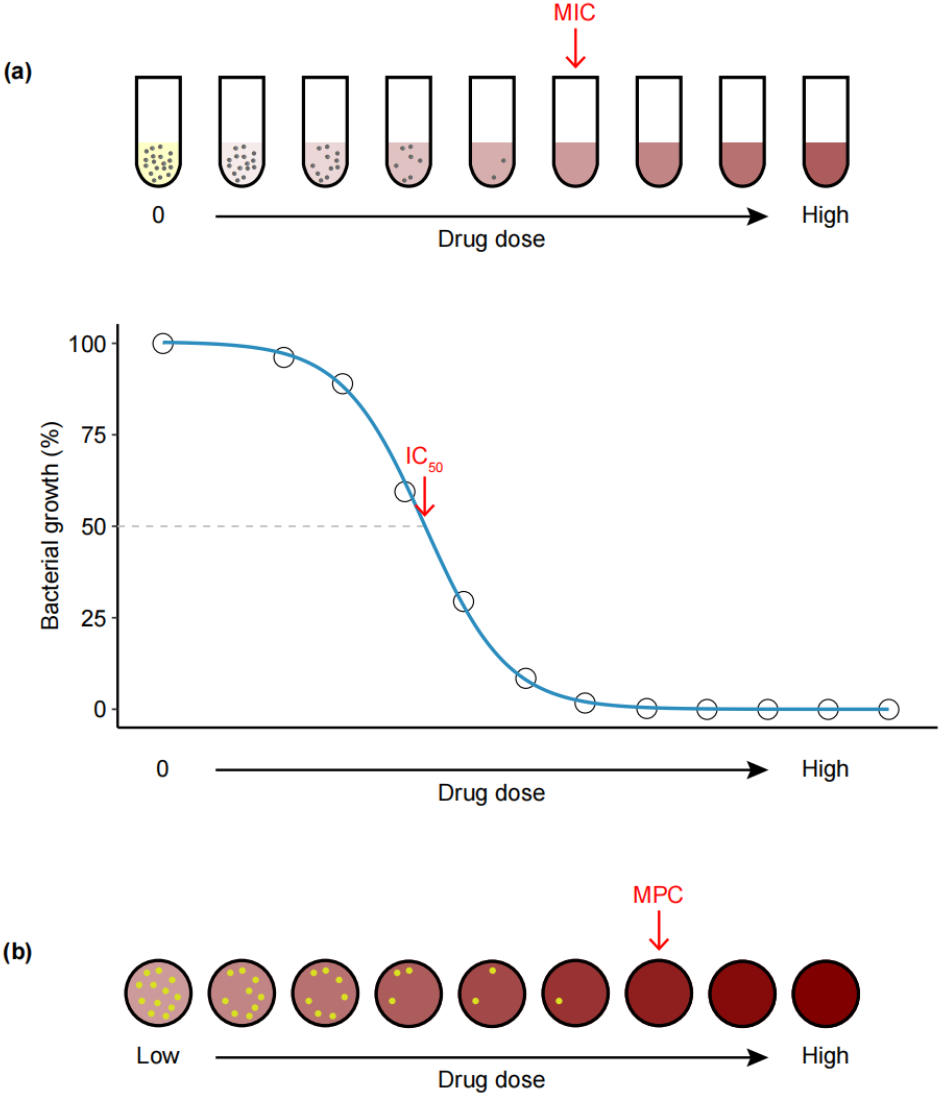
Schematic representation of measurement of minimum inhibitory concentration (MIC), half inhibitory concentration (IC_50_; a) and mutant prevention concentration (MPC; b) in antibacterial testing. Deeper reddish brown indicates higher drug dose supplemented into test tubes. The denser dots represent higher growth performance of bacterial population. Measures of MIC and IC_50_ involve growing the broth inoculated with bacterial cells along a gradient of antibiotic concentrations for a period of growth (usually < 24 h). The lowest drug concentration that blocks visible bacterial growth is recorded as MIC, and the function of dose-response curve is used to calculate IC_50_. Measure of MPC involves growing agar plates spread with bacterial cultures (typically with population size > 10^8^ CFUs) along a gradient of antibiotic concentrations; and the lowest drug concentration at which no colony grows is recorded as MPC.

An initially susceptible microbial population may evolve antibacterial resistance during the exposure, and survive drug treatments at concentrations higher than MIC. This is a well-known example of evolutionary rescue (Alexander et al., 2014; Andersson & Hughes, 2012; Bell, 2017). The efficacy of a drug in preventing resistance evolution may be measured using the mutant prevention concentration (MPC) index, defined as the lowest concentration at which a drug prevents the growth of the least susceptible, single-step, resistant mutant within a bacterial population (Fig. 1b) (Drlica, 2003; Zhao & Drlica, 2002). Higher MPC values indicate greater potential for a bacterial population to survive a drug via evolutionary adaptation.

Natural habitats are typically occupied by organisms of diverse species that form ecological communities, and community-level ecological functions may be maintained with dynamical species composition. This is particularly true for microbial communities of very high species diversity, as posited by the “it is the song, not the singer” hypothesis (Doolittle & Booth, 2016; Doolittle & Inkpen, 2018). Understanding how community-level functional properties respond to stress factors such as environmental pollution has been an important task (Dai et al., 2024; Griffiths & Philippot, 2013; Kandeler et al., 2000; Kuan et al., 2006; Shade et al., 2012). However, research in this filed has rarely adopted standardized measurement methods those are easily comparable across systems.

### Two measures of functional susceptibility of microbial communities to pollutants

Here we suggest adopting indices from antimicrobial susceptibility tests for microbial community research. First, the functional response of a community after immediate exposure to a gradient of stress can be used to estimate the resistance level, which may be measured using indices analogous to IC_50_ or MIC. Second, the community-level ecological functions may recover, to some extent, via species sorting (more resistant species becoming abundant) or evolutionary changes (more resistant genotypes increasing in frequency) under prolonged exposure, called community rescue (Bell et al., 2019; Fugère et al., 2020; Fussmann & Gonzalez, 2013; Low-Décarie et al., 2015). The potential for community rescue can be estimated from community-level functional response after a period of exposure allowing for community recovery, which is akin to MPC.

Therefore, two new indices are proposed here. First, the half inhibitory concentration (IC_50_), defined as the concentration at which a pollutant reduces a community-level function to 50% of the baseline state after immediate exposure (Fig. 2a). It is noteworthy that IC_50_ is different from the half maximal effective concentration (EC_50_) index commonly used in ecotoxicology research. The latter refers to the concentration of a drug that produces 50% of its maximum effect, and is often used as an estimate for the safety of drugs for animals (McClellan et al., 2008; Peters et al., 2013; Xiao et al., 2017; Yoshimura & Endoh, 2005). Second, the half community rescue prevention concentration (CRPC_50_), defined as the concentration at which a pollutant can prevent a community-level function from recovering to 50% of the baseline state during a period of prolonged exposure (Fig. 2b). Measurement of these two indices involves obtaining the dose-response relationship between the pollutant and a certain functional property of the microbial community. We performed two case studies to test this framework.

**Figure 2.**
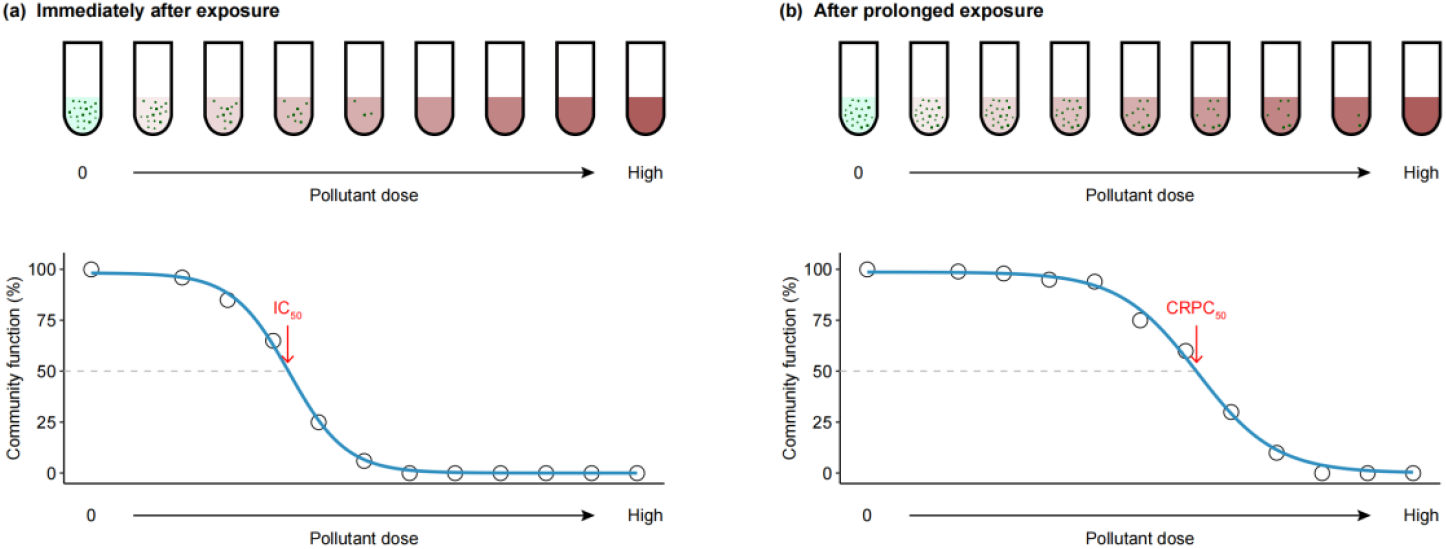
A graphical illustration of measurement of half inhibitory concentration (IC_50_; a) and half community rescue prevention concentration (CRPC_50_; b) of microbial communities. Deeper reddish brown indicates higher pollutant dose supplemented into the microcosms. The denser dots represent higher community function.

## MATERIALS AND METHODS

### Sample collection and preparation

Surface water samples collected from the North Moat, Beijing (116.38°E, 39.96°N) in July and September 2025 were used for Experiment I, and those collected from Wenyu River, Beijing (116.35°E, 40.13°N) in October 2025 were used for Experiment II. These original samples were filtered through nylon net (pore size 1 mm) for removing large organisms and particles. The filtered water sample was placed in a plastic container covered with nylon net (pore size 1 mm) in the open air for 1 d before subsequent assays.

Two pollutants, glyphosate and copper sulphate (CuSO_4_), were chosen in the present study. Glyphosate is the most extensively used herbicide in the past a few decades, with the chemical name *N*-(phosphonomethyl) glycine (Benbrook, 2016). It is widely used in the control of weeds in agriculture, gardens, land transportation systems, lakes and other environments. CuSO_4_ is used to prepare Bordeaux mixture, a time-honored fungicide which is widely used in vineyards and orchards (Deliopoulos et al., 2010), and also used as algicide and antimicrobial in water treatment (Liu et al., 2023; Watson et al., 2024). Large-scale use of these chemicals can cause toxic effects on non-target organisms including humans, a number of animals, plants and microbes (Bradberry et al., 2004; Briffa et al., 2020; Jeyaseelan et al., 2024).

### The experiments

Microcosms were set up by loading 2 L of water into each transparent cylindrical plastic bottle (30 cm of height, and 10 cm of diameter), covered with nylon net (pore size 1 mm). The microcosms were placed in plastic tanks filled with water of 15 cm of depth (for buffering daily temperature fluctuation).

In Experiment I, a gradient of glyphosate concentrations was made, a series of √2-fold dilutions from 1920 to 60 mg l^−1^, besides 0 mg l^−1^, with four replicates for each specific concentration (a total of 48 microcosms at each study season: 4 blocks × 12 microcosms). Samples were drawn from the microcosms at day 0 and 5 for functional assays.

In Experiment II, two pollutant agents were used. The glyphosate-treated microcosms covered the following gradient of concentrations: a series of √2-fold dilutions from 1920 to 60 mg l^−1^, and 0 mg l^−1^. Those under CuSO_4_ stress had a gradient of concentration of a series of √2-fold dilutions from 10.5 to 0.328 mg l^−1^ as well as 0 mg l^−1^. Two replicates were set up for each concentration. Samples were collected from the microcosms at day 0, 5 and 10 for functional assays.

### Measurement of community function

We considered respiration rate in darkness as a surrogate of community-level function, which is contributed to by every living member within a community. Water in each microcosm was mixed, 3 mL of which was transferred into a 3-mL sealed glass bottle with a PSt3 oxygen sensor spot at the bottom, by which the trajectory of dissolved oxygen (DO) concentration in the sample could be detected via SensorDish^®^ Reader (SDR) (PreSens, Germany). The bottles were set in darkness for avoiding oxygen generation from photosynthesis by photosynthetic microbes. The DO concentration of each sample was recorded every 1 min at room temperature for 24 h. We used the change rate of DO concentration from the 5th to the 20th hour to calculate respiration rate with the following exceptions. (a) If DO concentration at the 5th hour was higher than 250% air saturation (upper limit of SDR), we used the time point when DO concentration dropped to lower than 249% air saturation as the beginning point; (b) If DO concentration at the 20th hour was lower than 20% air saturation (growth of organisms might be restricted in extremely low oxygen environment), we used the time point when DO concentration dropped to lower than 20% air saturation as the end point.

### Statistical analysis

All statistical tests were carried out in the R environment (R Core Team, 2024). Log-Logistic model in “drm” function provided by R package “drc” (Ritz et al., 2015) was used to estimate IC_50_ and CRPC_50_. The average values of respiration rate at each pollutant concentration were used for the model fitting. Note that low concentrations of certain pollutants could enhance community growth, named “hormesis” (Calabrese & Baldwin, 2003; Southam & Ehrlich, 1943). This was true for our assays of cultures under glyphosate treatment. We excluded observations with respiration rates > 1.1-fold of the baseline state (0 mg l^−1^). IC_50_ and CRPC_50_ were compared using ratio tests wherein no differences are detected if the ratio contains 1 (Wheeler et al., 2006) by “comped” function provided by R package “drc”. *P* values were adjusted by Benjamini-Hochberg method (Benjamini & Hochberg, 1995) in Experiment II.

## RESULTS

### Experiment I

Immediately after exposure to glyphosate, microbial communities from the North Moat in July 2025 showed half inhibitory concentration (IC_50_) value of 83.88 ± 8.06 mg l^−1^ (Fig. 3a). After 5 days of exposure, the glyphosate concentration that could prevent microbial community respiration rate to recover to 50% level of the base line (no pollutant control) was 721.43 ± 98.30 mg l^−1^ (Fig. 3b); and the half community rescue prevention concentration (CRPC_50_) was roughly 9 folds of IC_50_ (*z* = 6.46, *P* < 0.001).

**Figure 3.**
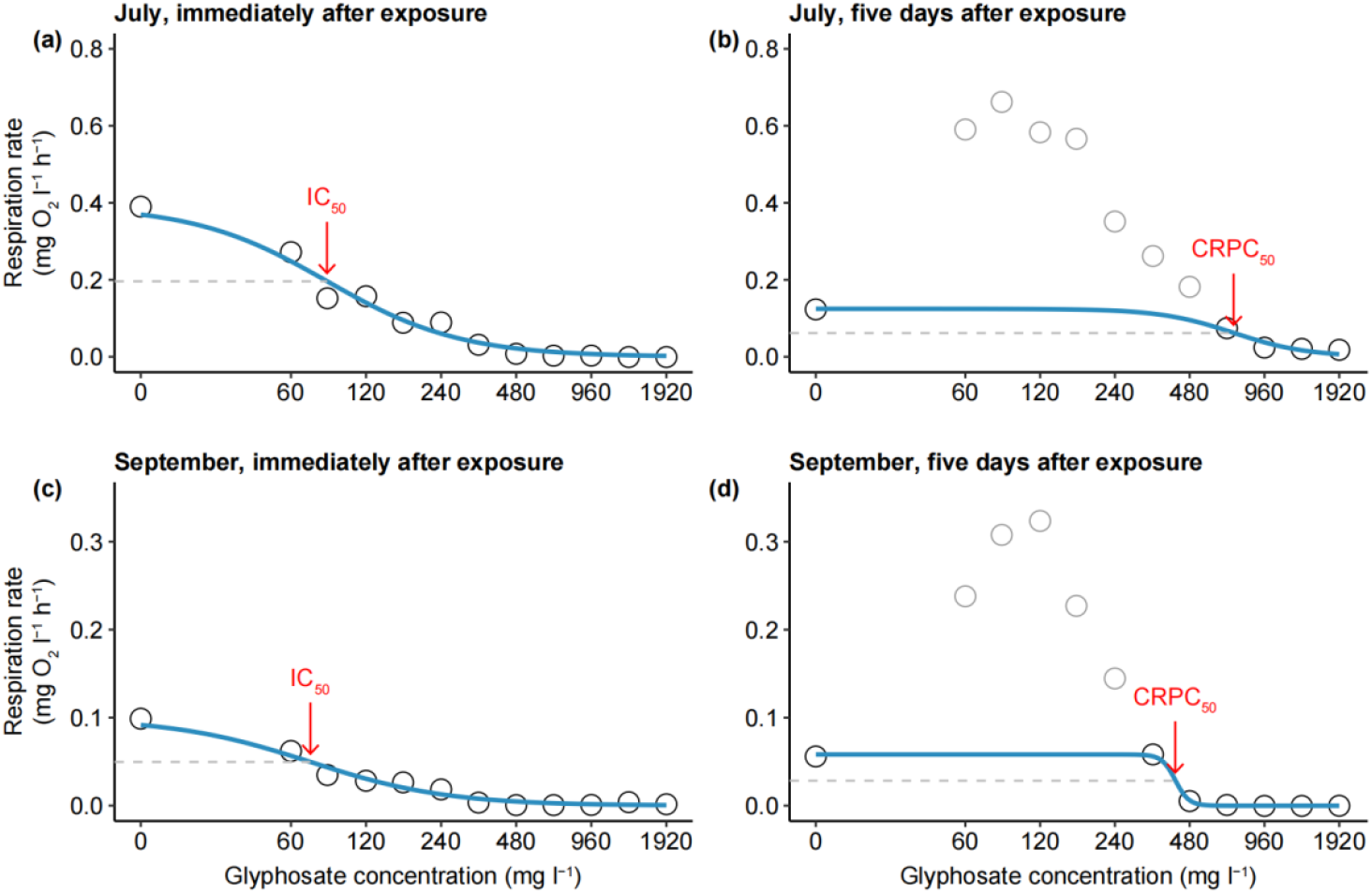
IC_50_ and CRPC_50_ showed by microbial communities in Experimental I. The blue line indicates the dose-response curve fitted by Log-Logistic model. The grey circles show those with obvious hormesis effect and were excluded in model fitting.

Microbial communities sampled in September 2025 showed IC_50_ value of 71.81 ± 12.55 mg l^−1^ (Fig. 3c), with no significant difference compared with those from July (*z* = 0.81, *P* = 0.418). The CRPC_50_ value was 419.97 ± 19.39 mg l^−1^ (Fig. 3d); this CRPC_50_ was roughly 6 folds of IC_50_ (*z* = 15.07, *P* < 0.001). The CRPC_50_ value measured in September was lower than that in July (*z* = 3.01, *P* = 0.003).

### Experiment II

Microbial communities collected from Wenyu River in October 2025 were treated with glyphosate or CuSO_4_. For glyphosate, the IC_50_ value (measured immediately after exposure) was 177.48 ± 25.07 mg l^−1^ (Fig. 4a). The CRPC_50_ values measured after 5 and 10 days of exposure were 649.96 ± 184.80 and 430.71 ± 105.93 mg l^−1^, respectively (Fig. 4b and c); and no significant difference was found between CRPC_50_ values measured at the two points in time (*z* = 1.03, *P*_adj_ = 0.303). The CRPC_50_ values both were significantly greater than IC_50_ (5 days, *z* = 2.53, *P*_adj_ = 0.030; 10 days, *z* = 2.33, *P*_adj_ = 0.030).

**Figure 4.**
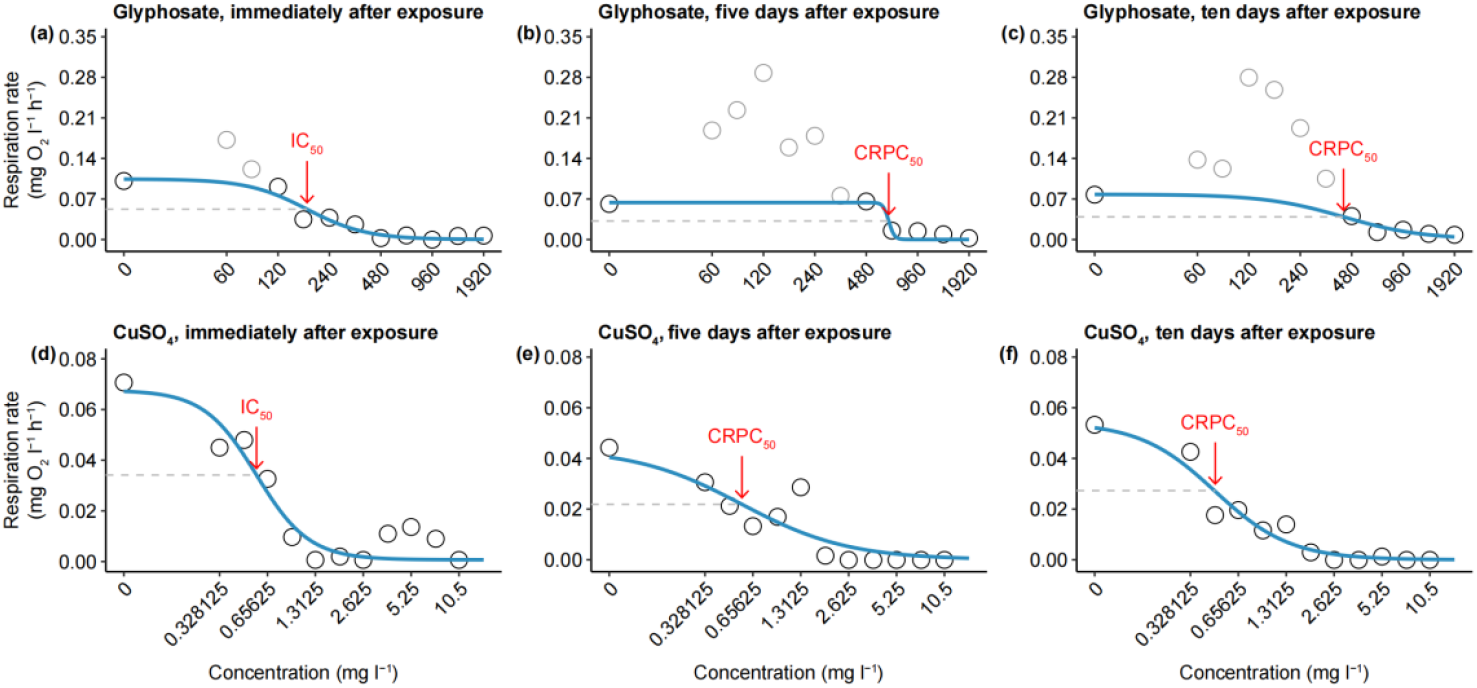
IC_50_ and CRPC_50_ showed by communities in Experimental II. The blue line indicates the dose-response curve fitted by Log-Logistic model. The grey circles show those with obvious hormesis effect and were excluded in model fitting.

For CuSO_4_, the IC_50_ value was 0.56 ± 0.08 mg l^−1^ (Fig. 4d), and the CRPC_50_ values measured after 5 and 10 days of exposure were 0.56 ± 0.15 and 0.47 ± 0.05 mg l^−1^, respectively (Fig. 4e and f). No significant difference was found between CRPC_50_ values measured at the two points in time (*z* = 0.55, *P*_adj_ = 0.871), both of which did not significantly differ from IC_50_ (5 days, *z* = 0.01, *P*_adj_ = 0.991; 10 days, *z* = 0.92, *P*_adj_ = 0.871).

## DISCUSSION

Microbial communities play crucial ecological roles in ecosystems. Assessing the community-level susceptibility is therefore essential for predicting ecological consequences of environmental pollutants. Here we introduce standardized measures for two key components of community-level susceptibility: functional resistance and community rescue potential. By adopting concepts from antibacterial testing, we propose two indices: the half inhibitory concentration (IC_50_) and the half community rescue prevention concentration (CRPC_50_). These measures would be comparable across environmental stress agents and study systems. In our demonstration assays, these two indices are obtained from the dose-response relationship between pollutant concentration and community respiration rate, a proxy for whole-community-level function.

### Assays of community properties

Assays of microbial community properties should take several issues into account. First, assays should be performed in a setting with as few changes as possible in environmental conditions compared with the natural habitats. It is recommended to load soil, sediment or water samples in experimental containers (microcosms) that are supplemented with only the tested pollutant (without nutrients). The microcosms should be incubated under conditions mimic the natural ones, ideally *in situ*.

Second, cautions are needed in deciding which ecological functions to study. A microbial community shows many functional properties (Escalas et al., 2019; Friedrich, 2011). Some functional processes are contributed to by a smaller portion of community members, e.g., photosynthesis is executed only by autotrophic organisms. Meanwhile, some functional properties reflect general metabolic activities of organisms; for example, respiration rate is contributed to by every living member within a community (Parker et al., 2016). Here we aim to measure the whole-community susceptibility, therefore, respiration rate serves as a proxy of whole-community function. Such functional process ensures that the measured response reflects the activity of all pollutant-resistant taxa within a community.

Third, the length of the exposure period should be taken in to account. Community rescue (recovery of community functional properties) take time to. With longer periods of pollutant exposure, there will be greater chances for the community to recover under higher doses of the pollutant. Meanwhile, change in species composition following stress exposure may incur the collapse in primary productivity in microcosms after an excessively long incubation time, resulting in decline of community function. Therefore, the CRPC_50_ value of a certain stress agent showed by a certain community may change with exposure period. Pilot assays should be carried out to figure out how long periods of exposure would be appropriate for particular pollutant agents.

### Model fitting

Organism growth often shows a monotonic decreasing trend with increasing pollutant dose, which can usually be described by logistic models (Ganie et al., 2017; Phillips et al., 2019; Seefeldt et al., 1995). However, some pollutants have hormesis effects on certain organisms, that is, low doses of pollutants enhance growth (Agathokleous et al., 2022; Erofeeva, 2022). There are two options to deal with this issue. First, the low doses that show hormesis effects may be excluded from analysis. Second, models including a parameter representing the hormesis effect can be used, as Brain-Cousens model and Cedergreen-Ritz-Streibig model (Brain & Cousens, 1989; Cedergreen et al., 2005). In our case studies, hormesis did not occur in all treatments. We removed observations with respiration rates > 1.1-fold of the baseline state from analysis and used a same model to fit dose-response curves of all treatments (Log-Logistic model).

### Demonstration experiments in the present study

Our Experiment I and II were performed to compare functional resistance and community rescue potential across study systems and environmental stress agents respectively. In Experiment I, communities from summer and autumn did not differ in functional resistance to glyphosate, but the summer community showed greater community rescue potential after prolonged exposure. One possible reason is that microbial communities from summer commonly harbour larger biomass and higher diversity due to warmer temperature or greater nutrient and radiance availability (Chen et al., 2025; Fang et al., 2023; Schnecker et al., 2023). Such communities with more complex structure are more likely to tolerate or recovery from environmental stress since there is more chance for them to contain resistant species or genotypes (Griffiths et al., 2003; Loreau, 2010).

In Experiment II, one particular community showed contrasting recovery dynamics under two different pollutant agents. Microcosms suffering glyphosate could recover function almost during the initial 5 days after exposure, whereas those suffering CuSO_4_ did not recover function even after 10-day exposure, suggesting that our microbial communities had less adaptive potential to the latter pollutant agent.

This finding explains why CuSO_4_ often shows long effective period as an algicidal. This chemical also has rather low market price (by weight). However, this also implies that copper ions residue resulted from agricultural and industrial activities may incur more severe and prolonged effects on the environment and public health.

Glyphosate can be degraded into harmless small molecules containing nitrogen and phosphorus by some microbes in the natural environment (Chen et al., 2022; Zhan et al., 2018). Such small molecules may provide nutrient substance for the growth and reproduction of various organisms and thus result in the hormesis as observed in our experiments and previous studies (Dabney & Patiño, 2018; Su et al., 2025).

In addition, it should be noted that some species in our study communities may perform anaerobic respiration and their respiration rates could not be represented by the decrease of DO. Nonetheless, for planktonic microorganisms in surface water that typically have weak anaerobic respiration, consumption of DO in the darkness has been the well-established approach to measuring respiration rates of communities (Cheung et al., 2024; Pace & Prairie, 2005); and studies of soil ecology also usually use CO_2_ flux as the proxy of respiration rates of microbial communities (Alster et al., 2023; Dove et al., 2021).

### Future perspectives

Empirical studies of community rescue are yet rare, though this concept has been proposed for over a decade (Fussmann & Gonzalez, 2013). The present framework can be a powerful and convenient tool for the development and improvement of community rescue theory. The measures of community susceptibility used in community ecology research may be concerned with changes in not only community-level properties such as biomass and net primary productivity but also species composition (de Grandpre & Bergeron, 1997; Fugère et al., 2020; Lepš & Lisner, 2025; Xu et al., 2025). The index which is measured based on community function in the present framework may also be measured based on species composition. For example, CRPC_50_ can be defined as the concentration at which a pollutant can cause 50% decline in species richness relative to the baseline state during a period of prolonged exposure.

## Notes

https://figshare.com/s/8bbb81d2f30a8f170d12

### Competing Interest Statement

The authors have declared no competing interest.

### Funding

- National Natural Science Foundation of China: 32371687

